# Single-nucleus co-expression networks of dopaminergic neurons support iron accumulation as a plausible explanation to their vulnerability in Parkinson’s disease

**DOI:** 10.1101/2022.12.13.514863

**Authors:** A. Gómez-Pascual, A. Martirosyan, K. Hebestreit, C. Mameffe, S. Poovathingal, T. G. Belgard, C. A. Altar, A. Kottick, M. Holt, V. Hanson-Smith, A. Cisterna, M. Mighdoll, R. Scannevin, S. Guelfi, J. A. Botía

## Abstract

**Motivation:** gene co-expression networks have been widely applied to identify critical genes and pathways for neurodegenerative diseases such as Parkinson’s and Alzheimer’s disease. Now, with the advent of single-cell RNA-sequencing, we have the opportunity to create cell-type specific gene co-expression networks. However, single-cell RNA-sequencing data is characterized by its sparsity, amongst some other issues raised by this new type of data.

**Results:** We present scCoExpNets, a framework for the discovery and analysis of cell-type specific gene coexpression networks (GCNs) from single-cell RNA-seq data. We propose a new strategy to address the problem of sparsity, named iterative pseudo-cell identification. It consists of adding the gene expression of pairs of cells that belong to the same individual and the same cell-type while the number of cells is over 200, thus creating multiple matrices and multiple scGCNs for the same cell-type, all of them seen as alternative and complementary views of the same phenomena. We applied this new tool on a snRNA-seq dataset human post-mortem substantia nigra pars compacta tissue of 13 controls and 14 Parkinson’s disease (PD) cases (18 males and 9 females) with 30-99 years. We show that one of the hypotheses that support the selective vulnerability of dopaminergic neurons in PD, the iron accumulation, is sustained in our dopaminergic neurons network models. Moreover, after successive pseudo-celluling iterations, the gene groups sustaining this hypothesis remain intact. At the same time, this pseudo-celulling strategy also allows us to discover genes whose grouping changes considerably throughout the iterations and provides new insights. Finally, since some of our models were correlated with diagnosis and age at the same time, we also developed our own framework to create covariate-specific GCNs, called CovCoExpNets. We applied this new software to our snRNA-seq dataset and we identified 11 age-specific genes and 5 diagnosis-specific genes which do not overlap.

**Availability and implementation:** The CoExpNets implementations are available as R packages: scCoExpNets for creating single-cell GCNs and CovCoExpNets for creating covariate-specific GCNs. Users can either download the development version via github https://github.com/aliciagp/scCoExpNets and https://github.com/aliciagp/CovCoExpNets

**Supplementary information:** supplementary data is available online.

## Introduction

Parkinson disease (PD) is the second-most common neurodegenerative disorder that affects 2-3% of the population ≥65 years of age^1^. In 2016, 6.1 million (95% uncertainty interval [UI] 5,0–7,3) individuals had Parkinson’s disease globally^2^, which projects to a staggering 12.9 million affected by 2040^3^. In most populations, 3-5% of PD is explained by rare variants identified in more than 20 genes, that is, representing monogenic PD^4^. On the other hand, 90 genetic risk variants collectively explain 16-36% of the heritable risk of non-monogenic PD^5^. PD diagnosis is clinically based. The Movement Disorder Society PD Criteria signals motor parkinsonism as the core feature of the disease, defined as bradykinesia plus rest tremor or rigidity. Lately, increasing recognition has been given to non-motor manifestations incorporated into both the current criteria and particularly into separate criteria for prodromal PD^6^. There is convincing evidence that the PD neurodegenerative process begins at least 20 years before the motor manifestations^7^, however, the accuracy of the clinical diagnosis of PD is still limited, especially in the early stages, when cardinal symptoms are not conclusive^8^. An accurate diagnosis of PD remains challenging and the characterisation of the earliest stages of the disease is ongoing.

The degeneration of midbrain dopaminergic neurons (DNs) within the substantia nigra pars compacta (SNpc) is a pathological hallmark of PD and Lewy body dementia^9^. However, not all dopamine-containing neurons at that region degenerate. This leads to the question of what are the molecular features that underlie selective neuronal vulnerability (SNV) of DNs in PD. Previous studies have suggested four different hypotheses to explain why DNs are more vulnerable in PD compared to other cell types. These are: i) dopamine can be toxic in certain conditions through the generation of reactive quinones during its oxidation; ii) iron is known to be able to generate reactive oxygen species (ROS) by the Fenton reaction and has been shown to accumulate with age at the SNpc; iii) pacemaking activity in SNpc DNs is accompanied by slow oscillations in intracellular calcium concentrations, causing extensive calcium entry and promoting chronically high levels of ROS production; iv) the massive scale of their axonal arborization, leads to very high numbers of axon terminals, elevated energetic requirements, and chronically high oxidant stress^10–12^.

In this paper, we look for evidence supporting any of these four SNV hypotheses using single-nucleus RNA-sequencing (snRNA-seq) data from human post-mortem SNpc tissue of 13 controls and 14 PD cases. From those we generate gene co-expression networks (GCNs) of specific cell types, to characterize the role of each cell type in conditions of normal and disrupted homeostasis (disease). GCN models identify gene co-expression/co-regulation patterns for the discovery of novel pathways and gene targets in both biological processes and complex human diseases^13^. Many methods have been proposed for constructing GCNs from both microarray and bulk RNA-Seq gene expression^14^, where weighted gene co-expression network analysis (WGCNA)^15^ has been widely used for the study of neurological diseases such as PD^13,16–19^, Alzheimer’s disease^20^, and Amyotrophic Lateral Sclerosis^21,22^. Bulk tissue datasets don’t have cell-type level granularity, but sc/snRNA-seq can give us the granularity we need to study the selective vulnerability at cell type level. Unfortunately, scRNA-seq comes with its own data hurdles, mainly high sparsity, i.e. large percentage of zeros, most of them being technical (dropouts events)^23^. In spite of that, GCN methods for bulk-based RNA-seq data can still be useful on sc/sn RNA-seq data. For example, WGCNA, perhaps the most widely used method for bulk expression data, has been applied to create sc/snRNA-seq GCNs^24–29^. On the other hand, some scRNA-seq specific methods address the sparsity problem. For example, scLink uses a filtering process to select only accurately measured read counts^30^. COTAN focuses directly on the distribution of zero UMI counts instead of focusing on positive counts^31^. HBFM generates latent factors to adjust the each cell’s gene expressions to alleviate from the zero-inflated and overdispersed attributes of scRNA-seq data^32^. Finally, scWGCNA uses aggregated expression profiles in place of potentially sparse single cells, where ‘metacells’ are constructed from specific cell populations by computing the mean expression from 50 neighboring cells using k-nearest neighbors^33^.

We propose a fast and multi-model method based on the notion of *pseudo cell*, i.e., new virtual cells created by adding, gene-wise, the counts of pairs of individual cells from the same type and individual. From the original matrix of n cells of a specific cell type, we get a new matrix of approximately n/2 pseudo cells aggregating all predecessor cells pairwise, within each donor. We repeat this while the size is still large enough. We create multiple expression matrices and the corresponding GCN for each. This implies managing multiple GCNs from the same experiment and cell type, which requires new ways of using the networks (i.e., new approach, new tools, and new methodology). While doing this, our assumption is simple: all GCNs obtained through this process may give us very robust findings (i.e. those that we repeatedly see across all GCNs) and alternative findings (i.e. those that emerge at one GCN and remain across the successors). Summing up all models’ new findings, we notably increase data insights while alleviating the sparsity bias. This approach has been implemented at the open source scCoExpNets R package and it has been applied mainly to the DNs population from our own cohort. In association with scCoExpNets, we have implemented the CovCoExpNets R package to help dissect subsets of genes within gene modules associated with more than one covariate (e.g., age and disease).

## Methods

### Post-mortem substantia nigra pars compacta samples used

The models we develop in this paper, both the cell-type GCNs and the covariate-specific models, are created using a discovery cohort of human PD cases and controls (N=27). To validate our results, we used a smaller, similar cohort (N=7).

The discovery cohort consists of post-mortem human SNpc tissues from 27 donors collected from the Oregon Health and Science University (OHSU), a total of 13 controls (8 males, 5 females) with 30-93 years and 14 cases diagnosed with PD (10 males, 4 females) with 57-99 years (supplementary Figure 1.a, b, c). RNA integrity number (RIN) ranges in 7.13 (6.93-7.33) and postmortem interval (PMI) in 20.94 (14.63-27.24) hours. The diagnosis (PD or control) was confirmed by a neuropathologist at OHSU after histological examination of the brain tissue. All postmortem brains were examined globally for plaques and tangles. Controls are non-neurological controls, meaning that they do not have any other observable neurological conditions at the time of death. All PD samples have Lewy body presence in midbrain and limbic brain areas while 44.4% also had it in the neocortical area. Amyloid plaque abundance was also evaluated for PD cases and represented in the scale 0-3, where 50% of the samples were classified at level 0, 42.86% at level 1 and 7.14% at level 2. Neurofibrillary tangles abundance was also evaluated for PD cases and represented in the scale 0-6, where 21.43% of the samples were classified at level 1, 14.29% at level 2, 21.43% at level 3 and 42.86% at level 4 (supplementary Figure 1.c.).

On the other hand, the replication dataset consists of post-mortem human SNpc tissues dissected from 7 samples provided by the Sepulveda brain bank. They are all male, 3 healthy controls with 70-88 years, and 4 PD donors with 61-79 years. RIN ranges in 7.13 (6.93 - 7.33) and PMI in 20.94 (14.63 - 27.24) hours.

### Single-nuclei RNA-sequencing

The tissue samples were processed at the Vlaams Instituut voor Biotechnologie (VIB), in Flanders, Belgium where the sample libraries were prepared. 2 mm biopsy punches were collected from the SNpc, nuclei were isolated and prepared, as per Mathys et al., 2019. The single-nuclei libraries were sequenced using Illumina HiSeq4000 10X Genomics Single Cell V2. The raw data were preprocessed using the 10× Genomics Cell Ranger 3.0.2.^35^. To quantify the unspliced mRNA present in the single nuclei RNA-Seq, we built a custom transcriptome using the GRCh38 reference that includes intronic regions. After generating single nuclei gene counts for each of the samples, we aggregated all samples with equalized read depth (via subsampling reads) to avoid batch effects introduced by different sequencing depths.

### Clustering and cell type identification

Data preprocessing for both discovery and replication purposes is the same. Raw UMI counts were generated by the Cellranger software. We used Seurat v3 R package^36^ to (1) remove MALAT1, which is known to be very highly expressed in single nuclei RNA-seq data and can drive the cell clustering, (2) to calculate the percentage of UMI counts coming from mitochondrially encoded genes for each cell and (3) filter out cells that have a unique feature (gene) count below 500 and above 6000 (this is the interval we defined to say that a gene is minimally expressed) and have a mitochondrial percentage (calculated in step 2) of above 5%.

Cell type identification was carried out following these steps: (1) perform SCT normalization for each sample separately (in this step, the mitochondrial percentage of each cell was also regressed out); (2) integrate the cells across all samples; (3) run PCA on integrated data; (4) run UMAP on a number of principal components; (5) cluster the cells based on principal components (supplementary Figure 1.d.); (6) exploring the expression of known marker genes, and finding differentially expressed genes (DEGs) between the cell clusters using the findMarkers function using the Wilcox method (supplementary Figure 1.e.). The Seurat R package in combination with an internal catalog of brain cell type markers was used to assign cell types to single nuclei. The genes used to identify cell types in the SNpc were SYT1 (neurons), GAD2 (GABA neurons), TH (DNs), SLC17A6 (glutamatergic, GLU, neurons), VCAN (oligodendrocyte progenitor cells, OPCs), DCN (connective tissue cells), MBP (oligodendrocytes), AQP4 (astrocytes), FLT1 (endothelial cells), CD74 (microglia) and THEMIS (T cells).

### Pseudo-cells approach to tackle sparsity: generating multiple scGCNs

We propose a new strategy to tackle sc/snRNA-seq data sparsity based on creating pseudo-cells. A pseudo-cell is the result of adding the gene expression of two cells that belong to the same individual and the same cell type cluster. We start from the initial gene expression matrix of a cell type and generate a GCN, denoted as T_0_ GCN. We create a new gene expression matrix in which each new pseudo-cell is created with pairs of cells of the same individual from the previous iteration (the new matrix has approximately half the cells than iteration T_0_) and create the corresponding GCN, denoted as T_1_ GCN. We repeat this process while the newly created expression matrix T_i_ (i > 1) has more than 200 pseudo-cells in size (Figure 1.a.). Let m be the minimum pseudo-cells to create a scGCN and n the number of T_0_ cells, we create up to log_2_(n/m) scGCNs.

**Fig. 1.**
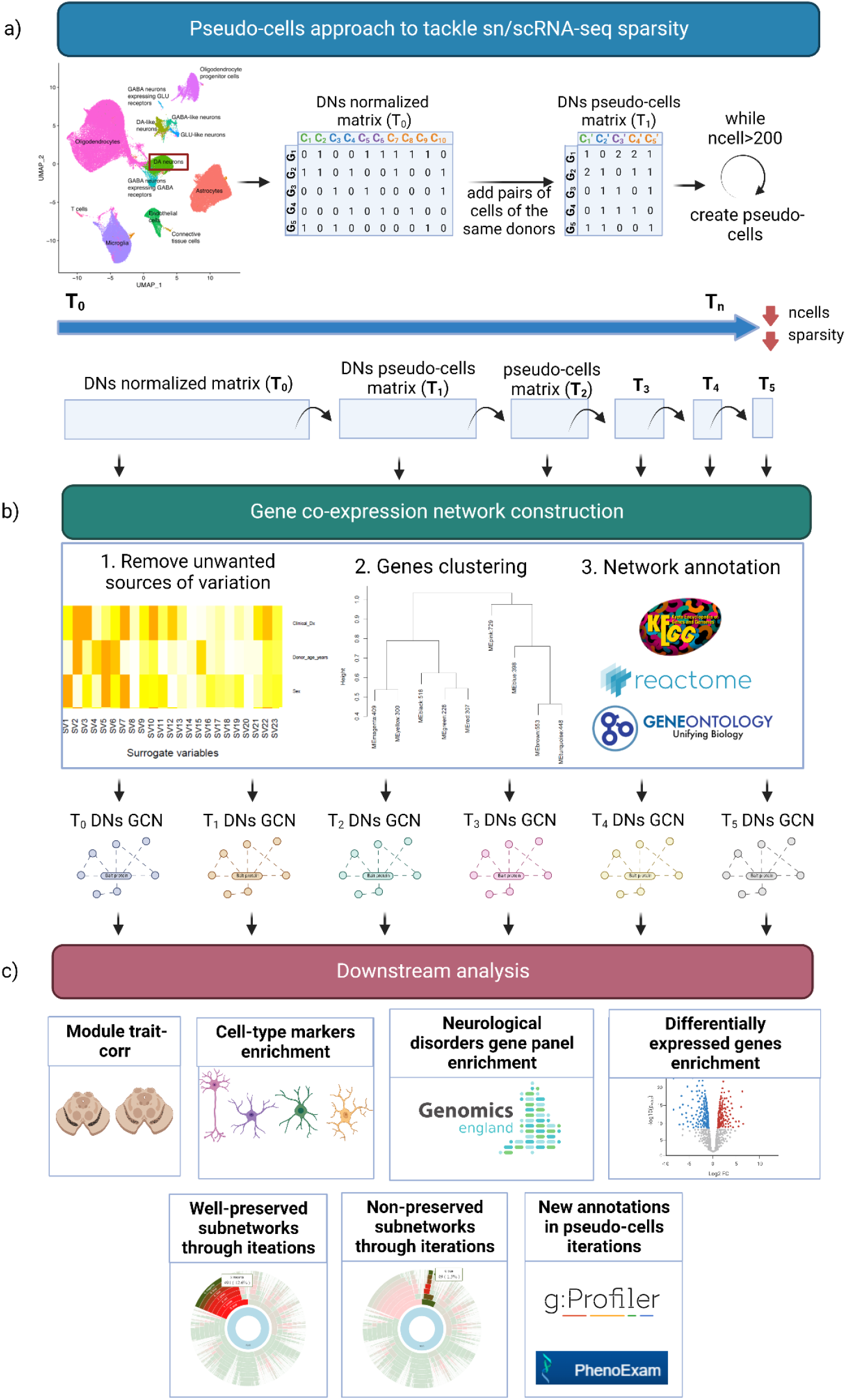
The creation and characterization of multiple scGCNs. **a)** Depicts the pseudo-cells approach of scCoExpNets R package to fight data sparsity. Gene expression cell pairs from the same cell type and individual are added to create a pseudo-cells expression matrix, significantly less sparse and with half the size than the predecessor. The scGCNs software repeats the process while the number of cells is still over 200. Each T_0_, T_1_ …, T_5_ as in the figure a) are used to generate a GCN. **b)** Creating the GCNs: technical covariates (i.e. RIN and PMI) are removed from gene expression using a surrogate variable analysis. Genes are clustered and each GCN module annotated using both functional (GO, REACTOME, KEGG) and phenotype (PhenoExam) enrichment analyses. **c)** Downstream analyses include identifying the most interesting gene modules by performing additional annotations on them (see plot).

### Gene co-expression network construction

We create a new GCN for each expression matrix T_0_, T_1_, T_2_ … For each T_i_, we followed the same three steps: data preprocessing, network construction and network annotation (Figure 1.b). Data preprocessing consists mainly of correcting for confounding effects of variables that are not of interest (technical and/or biological) with a surrogate variable analysis^37^. Specifically, our SNpc samples were corrected for RIN and PMI. Then, we created each network following the CoExpNets approach^38^, based on WGCNA. We first constructed a network *N* of gene-gene co-expression in the form of a squared *nxn* matrix, where *n* is the number of genes in the study and each *N*(*i, j*) is the interaction strength between the corresponding pair of genes (i.e. adjacency). Then, we used this as the basis for obtaining a new squared distance matrix with the distance between genes, ready to be used for obtaining clusters. These are built with a combination of hierarchical and k-means clustering. Finally, the modules of each GCN were annotated. First, we performed a functional enrichment analysis with gProfiler2^39^ including as databases the Gene Ontology (GO), the Kyoto Encyclopedia of Genes and Genomes (KEGG) and the Reactome Pathway database (REAC). The modules are also annotated using a phenotype enrichment analysis using the PhenoExam tool^40^, including as databases the Human Phenotype Ontology (HPO), Mouse Genome Informatics (MGI), CRISPRBrain, CTD, ClinGen, OrphaNET, UniProtm PsyGeNET, CGI and Genomics England. Finally, we added the module membership information for each gene in each module.

### Analysis of multiple scGCNs

We need the appropriate tools to simultaneously analyze a set of networks, derived from a set of pseudo-celluling expression matrices. We assume that all networks created through pseudo-cell technique may be potentially useful. And because the focus of the paper is studying SNV on DNs, we focus on networks built on DNs. And within the networks, we focus on specific modules of interest. These are modules mainly associated with PD and enriched for DNs markers (Figure 1.c.).

To identify modules associated with a sample trait, we checked if the module eigengenes (i.e. the first principal component of the expression profiles of all genes in a given module), were significantly correlated with the trait, e.g., clinical diagnosis, age.

To identify modules that are DN-specific or enriched for disease genes, we used a gene set enrichment analysis (GSEA) using the fgsea R package^41^. For such a purpose, we curated a list of gene sets as follows. Firstly, we included a list of PD genes, made up of PD genes from the latest PD GWAS^5^, PD genes with high or very high confidence obtained from the Blauwendraat et al., 2020 review and PD genes obtained from the Parkinson Disease and Complex Parkinsonism panel from Genomics England PanelApp. Secondly, we also included the *Neurology and neurodevelopmental disorders gene panel (level 2)*, made up of 28 gene panels, from the Genomics England PanelApp (accession in February 2022). We also included all cell type markers lists in the CoExpNets package. We added as well the differentially expressed genes (DEGs) at DNs for PD cases and controls detected by the Wilcox test as implemented in the *Seurat package*. Finally, we added our own manually curated sc/snRNA-seq DNs marker list based on the literature from the last 5 years (see supplementary table 1), including genes curated from 10 references^42–51^. This compounds a list of DNs markers in a variety of conditions, including species (human, mouse or even iPSc and models), brain areas (SNpc, VTA, midbrain) and inclusion criteria (clustering markers, based on literature, important for dopamine release…). Overall, we use 46 marker gene sets with variable sizes [10-1706] genes. We discarded not SNpc-specific gene sets and embryo and developmental stage markers.

In order to validate the modules of our GCNs, we apply a preservation test on an independent dataset, the Sepulveda dataset. The preservation test is implemented in the CoExpNets R package and is based on the WGCNA test for module preservation and generates a Z.press score. Under the null hypothesis of no module preservation, Z.press is below 2 while a Z summary greater than 5 (or 10) indicates moderate (strong) module preservation^52^. We use this same test to identify gene modules that, discovered at T_0_, remain throughout the successive iterations T_1_, T_2_…T_n_.

### Gene segregation based on covariates

Aging is a well known factor for PD^53–56^. Hence, it is common to detect gene modules associated with PD diagnosis and their age simultaneously (see results). We developed the CovCoExpNets R package to identify the genes that are exclusive to each association (but mainly to identify genes associated only with PD and not age) through the glmnet R package with LASSO optimization and create a covariate-specific GCN based on the genes detected (i.e. predictors with non-zero coefficients) (Figure 2, supplementary Figure 2).

**Fig. 2.**
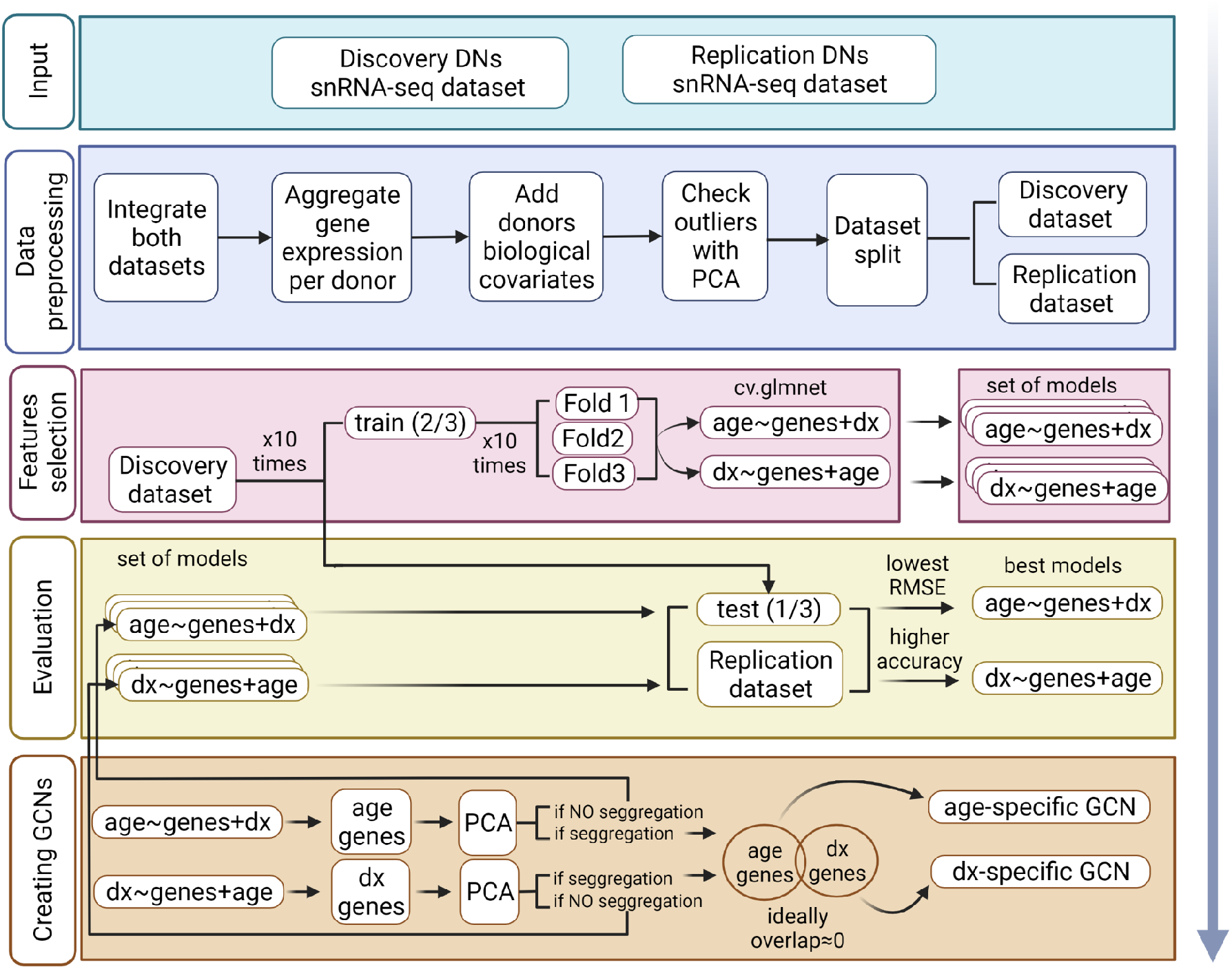
Overall pipeline for the creation of PD and age-specific GCNs from Oregon and Sepulveda datasets. The pipeline of the CovCoExpNets R package is described here. We follow a discovery and replication approach to generate networks on the discovery cohort (Oregon) and perform validation on the replication cohort (Sepulveda). We start with data preprocessing: we integrate discovery and replication datasets, and we aggregate the gene expression per donor into a single sample. Then we add age, diagnosis and sex as covariates to control for them and carry out a PCA at donor level to detect outliers and remove them if necessary. Once we do that, we split the dataset to separate donors into discovery and replication. Now the data is ready to select the best genes to be used as predictors for PD condition and age per donor. We call this *feature selection*: we split the discovery dataset into train (⅔ of the donors) and test (the other ⅓). We use the train data to build a model for age using as predictors the genes and controlling for diagnosis and sex using the cv.glmnet algorithm. We repeat the data sampling into train and test and model generation process 10 times. Within training, we split into three different folds for cross-validation. This is also repeated 10 times to alleviate variability of results. We perform the same process to build the PD model. The best model as selected by cross-validation is yet evaluated with the Sepulveda samples. For that, we test each model with both the test dataset and replication dataset and select the best model for each covariate. Finally, we create the covariate-specific GCNs: separately for PD and age models, we extract the selected predictors from the models (genes) and create a PCA based on the corresponding donor-level expression matrix to see if donors segregate per covariate. We create a module for each gene selected (see methods).

To create the age-specific GCN with CovCoExpNets (i.e. genes just associated with age but not PD), we obtained a LASSO model using as predictors, the individual-level expression matrix joined with the covariables of interest (i.e., clinical diagnosis and gender in this case) using age as the variable to model. We split the dataset into train (2/3 of the examples) and test (1/3) sets with 3-fold cross-validation within the cv.glmnet R function. Due to the small sample set, we repeated the train and test sampling 10 times, and the sampling for cross-validation, also 10 times. We then selected the best model based on the R^2^ on the validation set, made up of the remaining discovery samples (test) and the independent similar cohort from Sepulveda brain bank. We used predictors with non-zero coefficients as hub genes at the new GCN modules. Each hub gene generated a new module in the age-specific GCN. To complete the composition of each gene module with the rest of the genes, we created a basic linear model for the hub gene s

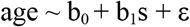

and kept adding genes to the model, choosing amongst the genes with highest correlation with the hub *s*. We do this while the R^2^ of the model improves. Each new module’s gene set was annotated with gProfiler2 with the sources GO, KEGG and REAC. We used a similar procedure to create the PD specific GCN.

## Results

### Cell-type composition and diversity in human SNpc

In the discovery cohort, 104,338 cells were identified, grouped into 14 major cell types and 24 annotated subclusters (supplementary Figure 1.d., supplementary table 2). We kept genes with reads within the range [500-6000]. The average observed read counts were 1817(1811-1821) 95% CI, where the highest values were detected in astrocytes/microglia 2640 (2531-2748) while the lowest values were detected in DNs 1329 (1316-1343) and GABA neurons expressing GABA receptors 1277 (1248-1306) 95% CI (supplementary Figure 1.f., supplementary table 2). The majority of the nuclei stem from oligodendrocytes (41.02%), followed by astrocytes (22.49%), microglia (14.51%) and oligodendrocyte progenitor cells (6.74%). Neuronal populations are relatively small in our samples, i.e. DNs are 6.29% of the total population (Table 3). The observed proportions in our data were consistent with other human SNpc studies. For example, (Agarwal et al., 2020) observe a proportion of 72% of oligodendrocytes in all glia they got from the SN of 5 healthy donors. (Wang et al 2022) detected 51.3% of oligodendrocytes, 13.1% of neurons, 9.4% of microglia and 8.4% of astrocytes from the SN of 23 idiopathic PD and 9 controls.

But we see a relatively high variability in cell proportion across samples, regardless of their condition (supplementary Figure 1.g.). Oligodendrocytes show the highest inter-sample variability, 13.94 (11.51-16.37)% for controls and 13.94 (11.91-15.97)% for PD cases while DNs show the lowest differences, 8.06 (6.64-9.48)% for controls and 5.34 (4.27-6.41)% for PD cases. Interestingly, we find statistically significant higher abundance of astrocytes in cases (Wilcoxon P<0.0193) and lower abundance of DNs-like neurons in the same group (Wilcoxon P<0.0107). Astrocytes protect neurons through the release of various neurotrophic factors but activated microglia can convert neuroprotective astrocytes into neurotoxic astrocytes ^57^. Previous studies demonstrated that A1 astrocytes - which lose the ability to promote neuronal survival, outgrowth, synaptogenesis and phagocytosis, and induce the death of neurons and oligodendrocytes - are abundant in various human neurodegenerative diseases including PD ^58^. This is consistent with what we see in our data.

### Networks are stable across pseudo-cell iterations

The original expression matrices of the different cell types (T0 expression matrices) have 77.22% (74.55-79.88) of zeros. We observed an association between sparsity and the number of detected genes (Pearson correlation −0.643, P<0.01307). We deal with such sparsity by creating new matrices of pseudo-cells named T_1_, T_2_, … with a minimum size of 200 pseudo-cells (see methods).

The average reduction in zeros in each cell type expression matrix from T_i_ to T_i+1_ is 14.84 (13.77-15.91)% (supplementary Figure 3). For each matrix, we created the corresponding GCN_0_, GCN_1_, GCN_2_, … within each cell type. A total of 112 scGCNs were generated, for 24 subclusters, grouped into 14 major cell-types. Networks were interpreted to be stable across pseudo-cell iterations when we observed structural features (i.e. number of modules, module average size), and biological features (i.e. number of functional enrichment terms per module) (see methods, supplementary Figure 3). We also observed that most of the modules at T0 were preserved across successive pseudo-cells iterations (supplementary Figure 3). A notable exception is GABA neurons expressing GABA receptors, where only 42.86% of the modules at T_0_ remain in T_1_, and only 33.33% of the modules at T_0_ remain in T_2_ and T_3_. A possible explanation for this exception is that this cell type presented the lowest number of genes detected (3150) but also one of the cell types with the highest number of modules detected in T_0_ (21 modules).

### The T_0_ dopaminergic neuron scGCN

Our main focus is to examine the evidence within our snRNA-seq data supporting any of the four selective vulnerability hypotheses of DNs in SNpc (see the introduction). Therefore, the rest of the manuscript focuses on the DN GCNs.

The DN scGCN at T_0_ is created from 6565 cells and 3890 genes. The network has 9 gene modules (432 genes per module on average). The criteria we used to identify interesting modules (see methods) is based on: (1) the association of the module eigengenes with clinical diagnosis and age; (2) enrichment on case/control DNs DEGs; (3) enrichment on cell type markers from both bulk and sc/snRNA-seq studies and (4) preservation of module structure in an independent dataset. Based on this, module M1 stood out as the most interesting module. It has 729 genes, it is correlated with clinical diagnosis (P<2.811·10^-218^), age (P<1.156·10^-244^) and sex (P<1.319· 10^-229^) (Figure 3a). It showed a significant GSEA enrichment (see methods) with case/control DNs DEGs (P<5.744·10^-7^). It als showed a significant GSEA for bulk RNA-seq DNs markers: the DNs markers collected from the CoExpNets R package (P<2.358·10^-3^) and the top 10% most specific SNpc genes for the bulk GTEx dataset (P<5.609·10^-5^) ^44^. M_1_ also presented the highest number of significant GSEA tests for sc/snRNA-seq DNs markers with a total of 18 significant tests, followed by M2 with 13 significant tests (Figure 3.b). This suggests M_1_ is DNs-specific. Moreover, the M_1_ module from T_0_ is preserved in the Sepulveda dataset with a Z summary pres > 2 (Z summary pres=3.393) (Figure 3c, supplementary table 3). Finally, M_1_ generated an abundant annotation term set from the functional enrichment analysis (438 terms, 23.06% of all terms) and from the phenotype enrichment analysis (74 terms, 18.00% of all terms).

**Fig. 3.**
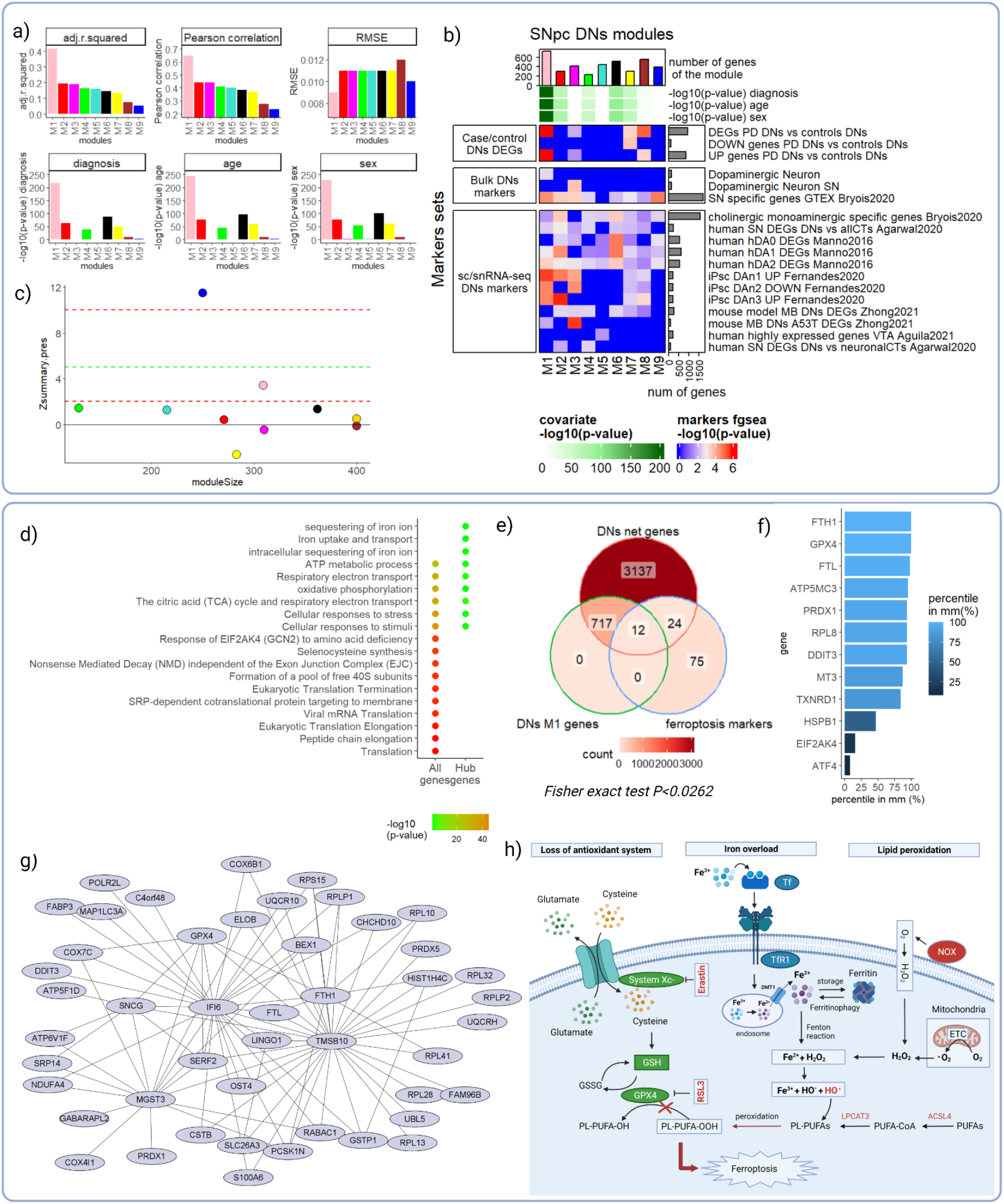
M_1_ module emerges as the most interesting SNpc DNs module. The top panel (Fig 3 a., b. and c.) gathers all evidence we use to select M_1_, a module of the T_0_ GCN of DNs, as the set of genes to focus on. **a)** Shows M1 (in pink) association with diagnosis, age and sex, in comparison to the other modules at T_0_ plotted with a variety of colors. RMSE refers to the rooted mean squared error between real and predicted values. **b)** Heat map with the significance level of annotation tests for each module (columns), M1 at the leftmost column (in pink). Bar height shows the number of genes per module. The green coloured heatmap under shows each module’ statistical significance of the association with clinical diagnosis, age and sex covariates, respectively. The rest of heat maps below show GSEA tests (see methods) with a variety of gene markers 1) DNs DEGs between PD cases and controls, 2) cell types markers from bulk RNA-seq studies obtained from CoExpNets R package and GTEx, and (3) DNs markers obtained from sc/snRNA-seq studies. Just marker gene sets, with at least one test as significant, shown. **c)** Preservation analysis of T_0_ DNs modules on the replication dataset (7 male cohort of PD cases and controls). M_1_ (pink) is more preserved than the rest (see methods), except done for blue. The bottom panel includes evidence linking M_1_ to the selective neuronal vulnerability hypothesis of iron accumulation. **d)** Enrichment plot showing the most significant annotations for M_1_ genes (left column) and the top 25 hub genes of M_1_ (right column); **e)** Venn Diagram between DN genes detected in our data, genes from M_1_ and ferroptosis markers from *FerrDb* database (Fisher exact test P<0.0262, see methods). **f)** Relevance of ferroptosis markers from FerrDb detected at M1 as measured in percentile of module membership (see methods). **g)** Top 50 genes M_1_ co-expression network. **h)** Ferroptosis pathway overview: loss of antioxidant system, iron overload and lipid peroxidation triggers ferroptosis.

### Evidence from M_1_ supports the iron accumulation hypothesis

Iron is highly abundant in the human body and a crucial component of hemoglobin. It is involved in the metabolism of catecholamine neurotransmitters and in the formation of myelin^59^. Selective accumulation of iron occurs in several brain regions and cell types in healthy aging and binds to ferritin and neuromelanin^60–62^. But abnormal accumulation of iron in specific brain regions occurs in many neurodegenerative diseases^63^. Different studies have shown that the accumulation of iron in PD patients is higher than in controls, especially in the SNpc and DNs^64–67^. Iron toxicity is due to its strong redox activity^68^. It easily reacts to H_2_O_2_ through the Fenton reaction producing neurotoxic intermediates or the end-products such as O_2_^-^ and OH·^69^. Since SNpc DNs are characterized by a high iron-content^64–67^, this leads to a higher production of Reactive Oxygen Species (ROS), piling up to the large amount of ROS produced in basal conditions which is necessary to support DN’s structure characterized by the massive arborization of their dendrites^70^. All this leads to an imbalance between ROS production and antioxidant defense inducing oxidative stress. This imbalance causes cell dysfunction and ultimately cell death^68^. Specifically, ROS contributes to ferroptosis, an iron-dependent non-apoptotic mode of cell death^71,72^.

A functional enrichment analysis on the 729 genes of M1 generated annotation terms like Parkinson disease (KEGG:05012, P<5.390·10^-34^), Ribosome (KEGG:03010, P<5.069·10^-62^), but also Oxidative phosphorylation (KEGG:00190, P<1.395·10^-36^), mitochondrial protein-containing complex (GO:0098798, P<3.380·10^-51^) (supplementary table 4). Interestingly, we repeated the functional enrichment analysis on the top 25 genes, i.e., the hub genes, based on their module membership. It turns out hub genes were enriched for Iron uptake and transport (REAC:R-HSA-917937, P<0.00274), Ferroptosis (KEGG:04216, P<5.612·10^-4^), Glutathione metabolism (KEGG:00480, P<0.01627), ferritin complex (GO:0070288, P<5.986·10^-5^) and intracellular sequestering of iron ion (GO:0006880, P<9.761·10^-4^) (see Figure 3d). To gain more insight about the possible connection of M_1_ with iron metabolism and ferroptosis, we crossed its genes with a list of 111 ferroptosis markers obtained from *FerrDb*, a manually curated resource for regulators and markers of ferroptosis^73^. We found 24/111 markers as detected genes in our DNs dataset (Figure 3e.). Half of them were found at M_1_ (12/24) (Fisher Exact Test P<0.0262), where 58.33% (7/12) of them showed a percentile in MM over 90%. The most relevant genes included Ferritin heavy chain (FTH1) and Glutathione peroxidase 4 (GPX4) (Figure 3f and 3g). GPX4 plays a pivotal role in the occurrence of ferroptosis, it converts GSH into oxidized glutathione and reduces the cytotoxic lipid peroxides (L-OOH) to the corresponding alcohols (L-OH). Inhibition of GPX4 activity can lead to the accumulation of lipid peroxides, a hallmark of ferroptosis^72,74^. GPX4 is down-regulated in our case/controls DEGs (P<4.647·10^-13^, log2FC= −0.182). All this evidence may suggest that ferroptosis is disrupted in PD DNs (Figure 3h.).

### Evidence of M_1_ as a well-preserved module across iterations

We observed that module M_1_ is stable across successive pseudo-cell matrices and scGCNs, suggesting that M_1_ is a transcriptional signal robust to the underlying data rather than simply an artifact. Supplementary Figure 4a. includes a visual representation of this by means of a sunburst plot^75^. The inner ring shows M_1_ at T_0_, composed of 729 genes, the largest in this network. M_1_ at T_1_ keeps 598 of those genes, 595 at T_2_, and so on until it reaches 491 at T_6_. Therefore, this module is robust against data sparsity. In consequence, the features of these modules in terms of cell type markers enrichment and case/control DNs DEGs enrichment is quite similar through pseudo-cells iterations (see supplementary Figure 4c.). In addition, M_1_ module from T_0_ is preserved in the Sepulveda dataset in most of the iterations with a Z summary press of 2.497 (1.955 - 3.040) (only T_1_ Z summary press is slightly under the cutoff).

### Non-preserved modules provide new insights

Some modules at T_0_ DNs GCN changed their gene composition considerably, throughout the pseudo-cell iterations. Supplementary Figure 4b. shows the example of a module which is made up of 300 genes in T_0_ but, after pseudo-cells iterations, only 49 of these genes remain grouped together in the last iteration, while the rest of the genes are regrouped into other modules. This has the potential of transforming some of them into newly relevant modules. For the module highlighted at at that figure, we can see how its association with disease gets sharper across iterations together with its association with age and sex. We also see how it improves its enrichment in both case/control DNs DEGs and DNs markers (supplementary Figure 4d.). Another interesting phenomena emerged when we studied enrichment of the 26 PD-related HPO terms (OMIM: 168600) (see supplementary table 5) across pseudo-cell iterations in this same module. Interestingly, movement related terms like rigidity (HP:0002063), tremor (HP:0001337) or parkinsonism (HP:0001300) became less significant across pseudo-cell iterations. Notably, this includes iterations where the correlation with the diagnosis is the highest (T_2_ and T_3_). Just the opposite happened with HPO terms like mask-like facies (HP:0000298), dysarthria (HP:0001260) or dystonia (HP:0001332). This suggests that pseudo-cell iterations have the potential to specialize modules into more specific aspects of PD. Actually, we found that 26.92% of the HPO terms are just found on DNs GCNs other than that of T_0_. They include dysarthria (HP:0001260), hallucinations (HP:0000738), mask-like facies (HP:0000298), sleep disturbance (HP:0002360), sporadic (HP:0003745) and urinary urgency (HP:0000012).

### Segregating gene-level associations across relevant covariates

M_1_ from the T_0_ scGCN emerged as the most interesting module at T_0_. This module showed associations with clinical diagnosis, age and sex (Figure 3a). Conventional co-expression analyses would assign all genes within M1 to all three covariates. We wanted to disentangle these associations mainly to identify genes in M1 just associated with PD, and genes just associated with age. For such a purpose, we designed the concept of **covariate-specific gene co-expression networks** (see methods) and created a DNs PD-specific GCN and an age-specific GCN. We started by integrating the DNs expression matrices from the discovery cohort (Oregon dataset) and the replication cohort (Sepulveda dataset) into a single expression matrix to normalize genes across samples. The UMAP plot in Figure 4a. shows both cohorts integrated across the space. By collapsing cells into donors by averaging gene expression per individual, we created a donor-level expression matrix, with the 35 donors (28 Oregon samples and 7 Sepulveda samples) and 2529 genes. A total of 3 donors emerged as outliers in a PCA plot (Figure 4b). Amongst the outliers, one of the samples had the highest PMI (61 h) and, accordingly, the lowest RIN (6.4). A second sample was just 30 years old (medium age is 78 years). And all three donors had less than 5 DNs each. After removing them from the dataset, we repeated the PCA analysis and found no undesired association with any covariate (see Figure 4c.).

**Fig. 4.**
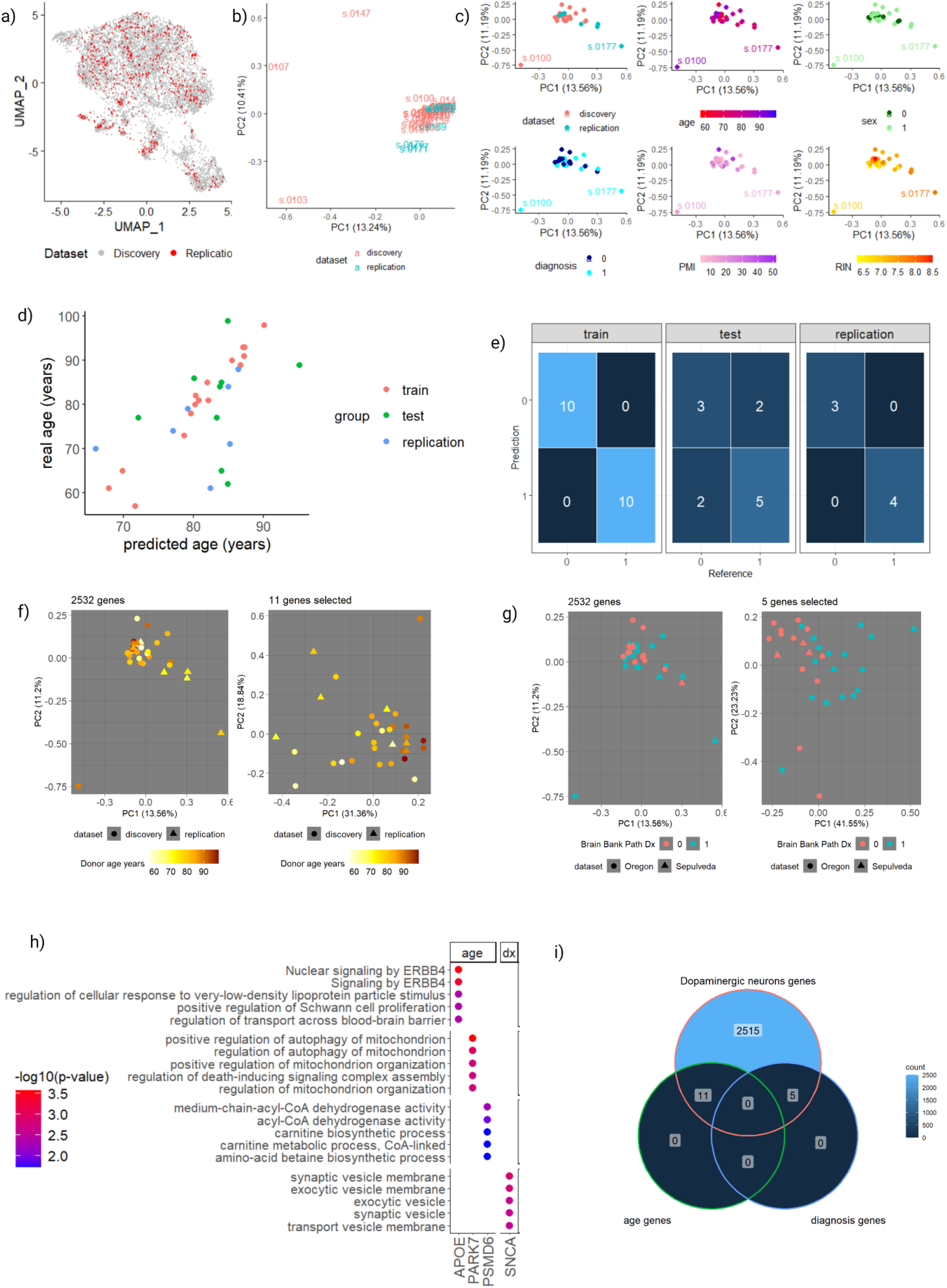
Segregating donors per covariate. **a)** UMAP projection of discovery and replication datasets integrated at single-cell level; **b)** PCA projection at donors level including all the donors colored by dataset (discovery or replication); **c)** PCA projection at donors level after removing the outliers (s.0147, s.0107, s.0103) colored by different criteria (dataset, age, sex, diagnosis, PMI, RIN); **d)** real vs predicted age for each individual is represented. Donors are colored by the set they belong to (train, test or replication set). **e)** confusion matrix for the discovery dataset (train and test) and the replication dataset (the lower diagonal represents the hits); **f)** PCA projection at donors level, where donors are colored by their age and the shape represents the dataset where they belong. In the picture on the left, all the features were used and we see no segregation per age. In the picture on the right, we only used the 11 age-specific genes selected and, as we move from left to right, we see donors from the youngest to the oldest. **g)** PCA projection at donors level, where donors are colored by their age and the shape represents the dataset where they belong. In the picture on the left, all the features were used and we see no segregation per diagnosis. In the picture on the right, we only used the 5 PD-specific genes selected and donors are segregated per diagnosis. **h)** top 5 most relevant annotations for the age-specific and diagnosis-specific most annotated modules. x axis represents the leader of the module, y axis represents the term name and the color represents the -log_10_(p-value). **i)** venn diagram represents the absence of overlap between age-specific genes and diagnosis-specific genes selected with CovCoExpNets.

We then used the CovCoExpNets R package to generate models of age from the training samples. The best model for predicting age had 0.765 R^2^ on the training set. Figure 4d. shows predicted vs observed age in years for each individual, grouped by set (train, test or replication set). We then plotted all donors in a PCA space using only the 11 age-specific genes detected by CovCoExpNets. The 1st PCA showed a Pearson correlation with age of 0.565 (P<7.637·10^-4^) using all the samples (Figure 4f.). The selected genes for predicting age are used as hub genes for initially empty modules. At the end, 28 genes (11 hubs plus 17 new genes) made up the age-GCN (supplementary table 6). We found a total of 1404 annotations, where 37.61% of them belong to the PARK7 module, 34.40% to the APOE module and 11.04% to the PSMD6 module (supplementary table 7).

ApoE participates in the distribution/redistribution of lipids among various tissues and cells of the body by serving as the ligand for binding to specific cell-surface receptors^76^. A relationship between aging and the APOE-4 allele has been previously described. Honea et al., 2009 demonstrated that, in nondemented older adults with the E4 allele, cognitive performance was reduced, and atrophy was present in the hippocampus and amygdala compared to APOE4 negative participants. Moreover, Y. J. Li et al., 2004 demonstrated that the APOE-4 allele increases risk and decreases age at onset of PD, an association that may not be dependent upon cognitive impairment. The APOE module is implicated in signaling by ERBB4, APOE gene transcription is stimulated by ERBB4s80^79^, and regulation of transport across the blood-brain barrier (see Figure 4.h.).

On the other hand, Thomas et al., 2011 demonstrated that loss of DJ-1 (PARK7) leads to loss of mitochondrial polarization, fragmentation of mitochondria and accumulation of markers of autophagy (LC3 punctae and lipidation) around mitochondria in human dopaminergic cells. These effects are due to endogenous oxidative stress, as antioxidants will reverse all of them. In this regard, the most relevant annotations of the PARK7 module are implicated in the autophagy of mitochondria (see Figure 4.h). Initially, Van Duijn et al., 2001 identified PARK7 as a novel locus for autosomal recessive Early-Onset Parkinsonism. More recently, Gialluisi et al., 2021 identified 28 disrupting variants in 26 candidate genes for late-onset PD, where PARK7 gene was also included, suggesting that PARK7 mutations are not exclusive to early onset forms of PD. In addition, they noticed that, in several families, single mutations in recessive genes such as PARK2, PINK1, PARK7, were co-inherited with variants in other candidate genes, suggesting that they might play a role as risk factors in the heterozygous status.

In regard to the model of PD, the best model for predicting diagnosis correctly classified all controls and all PD cases of our train set. Furthermore, this model correctly classified most controls (3/5) as well as most PD cases (5/7). And even more interesting is that this model correctly classified all the samples coming from the replication set. All the confusion matrices are represented in Figure 4e. We also demonstrated that we are able to segregate the 32 donors per clinical diagnosis in a PCA space using only the 5 diagnosis-specific genes (Figure 4g.). Donors from the replication dataset follow this same trend. Then, we created the diagnosis-GCN, made up of 12 genes. We found a total of 676 annotations, where 78.85% of these annotations belong to the SNCA module. SNCA is considered as the major causative gene involved in the early onset of familial Parkinson’s disease (FPD) characterized by five missense mutations identified so far^83^. The normal function of α-synuclein remains enigmatic, despite more than 25 years of research, however, α-synuclein’s presynaptic localization and its interaction with highly curved membranes and synaptic proteins strongly suggests a regulatory function associated with the synapse^84^. In this regard, the top most relevant annotations of this module are associated with synaptic vesicle membranes (see Figure 4.h.).

Interestingly, we found no overlap between the 11 age-specific genes and the 5 diagnosis-specific genes, that is, they are specific to each covariate (Figure 4.i).

## Discussion

SnRNA-seq is just starting to show all its possibilities as a means to enable bioinformatic analyses on the right cell type in the right space and time^85,86^. One of the main challenges of snRNA-seq data is the sparsity of gene expression matrices^23^, e.g, we find around 75% of zeros, on average, in our expression matrices. Therefore, we developed the scCoExpNets R package with this in mind. It is focused on using single-cell gene expression’s natural sparsity to our own advantage. scCoExpNets reduces sparsity through pseudo-cells, i.e. pairwise aggregation of cells within the same cell type and donor. By using this technique, we introduce the idea of multiple models through the creation of successive network models across T_0_, T_1_, …, T_n_ expression matrices. Thanks to this we obtain transcriptomic signals that are robust to sparsity versus signals that are artificial, increasing our trust in them and also modules which evolve towards potentially more interesting gene groups.

We applied scCoExpNets on a snRNA-seq dataset human post-mortem SNpc tissue of 13 controls and 14 PD cases (18 males and 9 females) with an age range of 30 to 99 years. All the samples in the discovery cohort yielded a cell population of 104338 cells, segregated into 14 main cell types. Out of all these cells, we generated and studied models for all the cell types. In this paper, we focused on the population of DNs as we were looking for evidence to shed light on the selective vulnerability in that cell type. The M_1_ model from iteration T_0_ emerged as the most interesting of all DN modules. This module is preserved in an independent cohort of 7 males, which makes it more reliable. Moreover, the gene composition of this module is maintained through the pseudo-cells iterations, therefore, it is quite robust to sparsity. All the evidence emerging from the analyses we have performed on M_1_ points towards suggesting that the weakest point in DNs within PD cases lies on the dysregulation of iron metabolism.

In the brain, iron is involved in many fundamental biological processes including oxygen transportation, DNA synthesis, mitochondrial respiration, myelin synthesis, and neurotransmitter synthesis and metabolism^63^. In healthy ageing, selective accumulation of iron occurs in several brain regions and cell types, with iron mainly bound within ferritin and neuromelanin. Total iron concentrations increase with age in the SNpc, putamen, globus pallidus, caudate nucleus, and cortices. Moreover, in the SNpc, ferritin heavy chain (FTH1) and ferritin light chain (FTL) concentrations increase with age, whereas these are constant in the locus coeruleus; therefore iron could contribute to neurodegeneration in the substantia nigra more than in the locus coeruleus^63^. Interestingly, we have detected FTH1 and FTL as hub genes in the M_1_ module. When iron levels exceed the cellular iron sequestration capacity of storage proteins or other molecules, iron is loosely ligated (“labile”) and able to participate in redox reactions collectively referred as “Fenton chemistry”, a series of reactions where labile iron reacts with endogenously produced H_2_O_2_ or O_2_^-^ to form oxygen radicals (i.e. hydroxyl and peroxyl)^87^, both able to generate a lipid peroxide (ROOH)^88^. Lipid peroxidation is the key downstream feature of ferroptosis, a non-apoptotic, iron dependent form of regulated cell death^74^. The key regulators of lipid peroxides in cells are the glutathione peroxidase (GPx) enzymes, particularly GPx4, which uses GSH as a co-substrate to reduce lipid peroxides to the corresponding alcohols, therefore, inactivation of this enzyme results in the accumulation of lipid peroxides and often cell death^88^. In this regard, the GPX4 gene has been detected as a hub gene in the M1 module. Dixon et al., 2012 also identified a distinct set of genes that regulate the ferroptotic mechanism, including ribosomal protein L8 (RPL8), iron response element binding protein 2 (IREB2), and ATP synthase F_0_ complex subunit C3 (ATP5G3). In the M_1_ module, we have detected RPL8 and ATP5MC3 as hub genes (ATP5G3 is the previous symbol for ATP5MC3 gene). Finally, Do Van et al., 2016 demonstrated that ferroptosis is an important cell death pathway for DNs in PD. In addition, treatments that protect against ferroptosis have shown therapeutic potential in PD^90,91^.

On the other hand, the CovCoExpNets, in association with scCoExpNets, helps in dissecting the nature of gene modules associations with more than one covariate or factor (e.g., age and disease). In this paper, it is particularly useful to select genes exclusively associated with a condition, e.g., PD, or age. For that, we create covariate-specific gene-co-expression networks. We believe that many co-expression based studies can benefit from using this new type of analysis when network modules are so complex and need breaking down into simpler and more specific gene sets.

To this end, we have applied the LASSO regularization mechanism to detect the age-specific genes and the diagnosis-specific genes separately on the training set (2/3 of discovery dataset) and we have tested the predictive capacity of these models in the test and the replication sets. In this regard, we have identified 11 age-specific genes and 5 diagnosis-specific genes that do not overlap and can be used to segregate the donors in a PCA space. We have used these hub genes to create the corresponding age-specific GCN and diagnosis-specific GCN, respectively. We have detected that the age-specific modules led by PARK7, APOE and PSMD6 are the most annotated ones. The same happens for the diagnosis-specific module led by SNCA.

Interestingly, we found no overlap between the 11 age-specific genes and the 5 diagnosis-specific genes, suggesting that we were able to differentiate age-specific and PD-specific signals in our gene models. For example, initially, PARK7 was identified as a causal gene for early-onset PD but we have identified it as age-specific, therefore, which would lead to future avenues of research regarding the role of this gene in late-onset PD. More research is needed to develop a strategy to ensure that we are controlling for the age when predicting diagnosis and vice versa.

## Supporting information

Supplementary table 1. Curated list of dopaminergic neurons markers.

Supplementary table 2. Single-nucleus discovery dataset main statistics.

Supplementary table 3. Dopaminergic neurons T0 modules preservation.

Supplementary table 4. Dopaminergic neurons T0 M1 module enrichment.

Supplementary table 5. HPO terms of Parkinson disease, late onset (OMIM:168600).

Supplementary table 6. Dopaminergic neurons age-specific network structure.

Supplementary table 7. Dopaminergic neurons age-specific network enrichment.

## Code availability

The scCoExpNets R package is available at https://github.com/aliciagp/scCoExpNets

The CovCoExpNets R package is available at https://github.com/aliciagp/CovCoExpNets

---

**Supplementary fig. 1.**
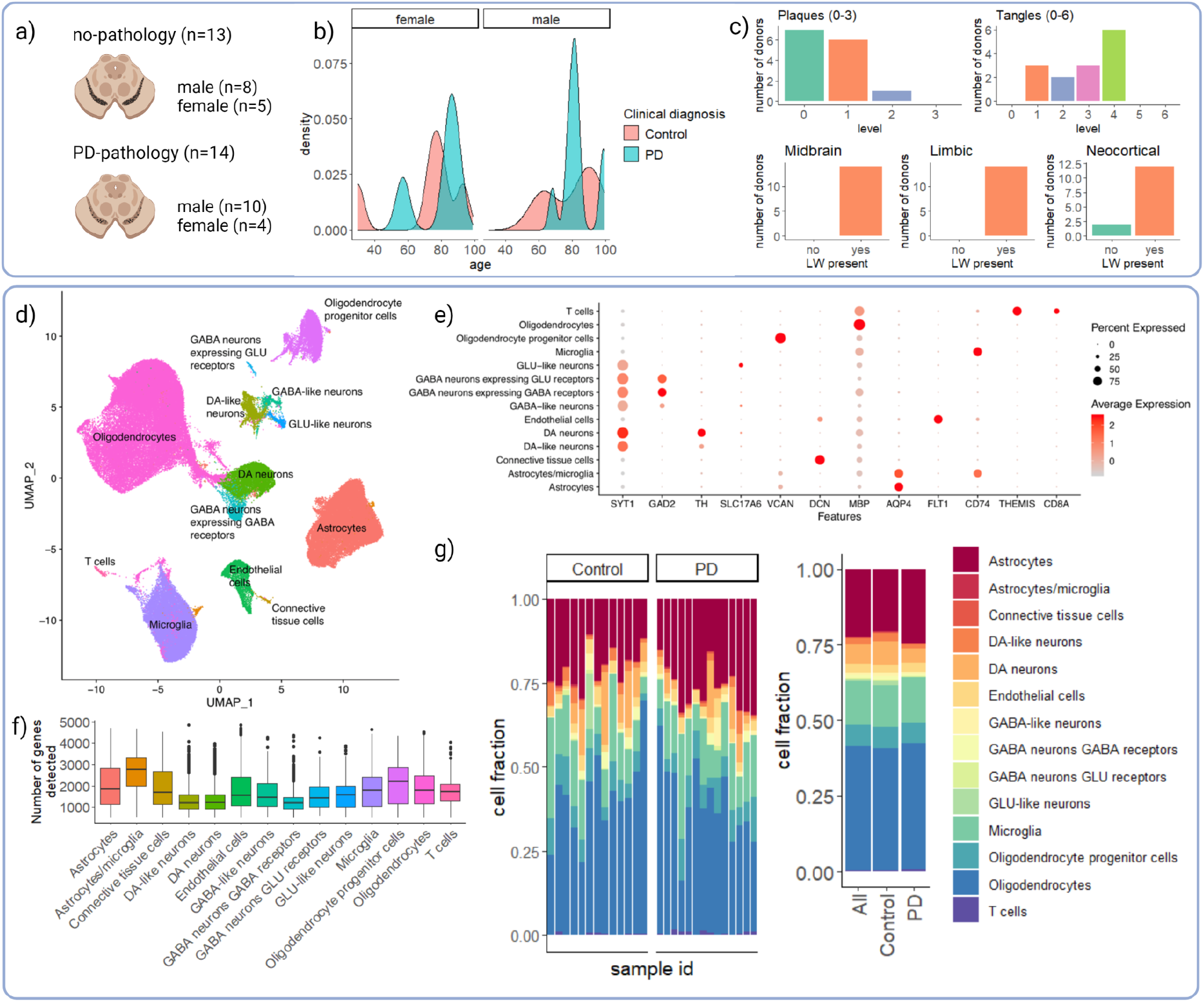
The substantia nigra single-nucleus RNA-seq dataset. **a)** Number of donors per diagnosis and sex; **b)** age distribution per clinical diagnosis and sex; **c)** PD cases plaques and tangles classification and lewy body presence in midbrain, limbic and neocortical areas; **d)** UMAP projection of cells annotated per cell type; **e)** average expression of key cell specific markers across cell types used; **f)** distribution of the number of genes detected per cell type; **g)** cell type proportion per sample and clinical diagnosis.

**Supplementary fig. 2.**
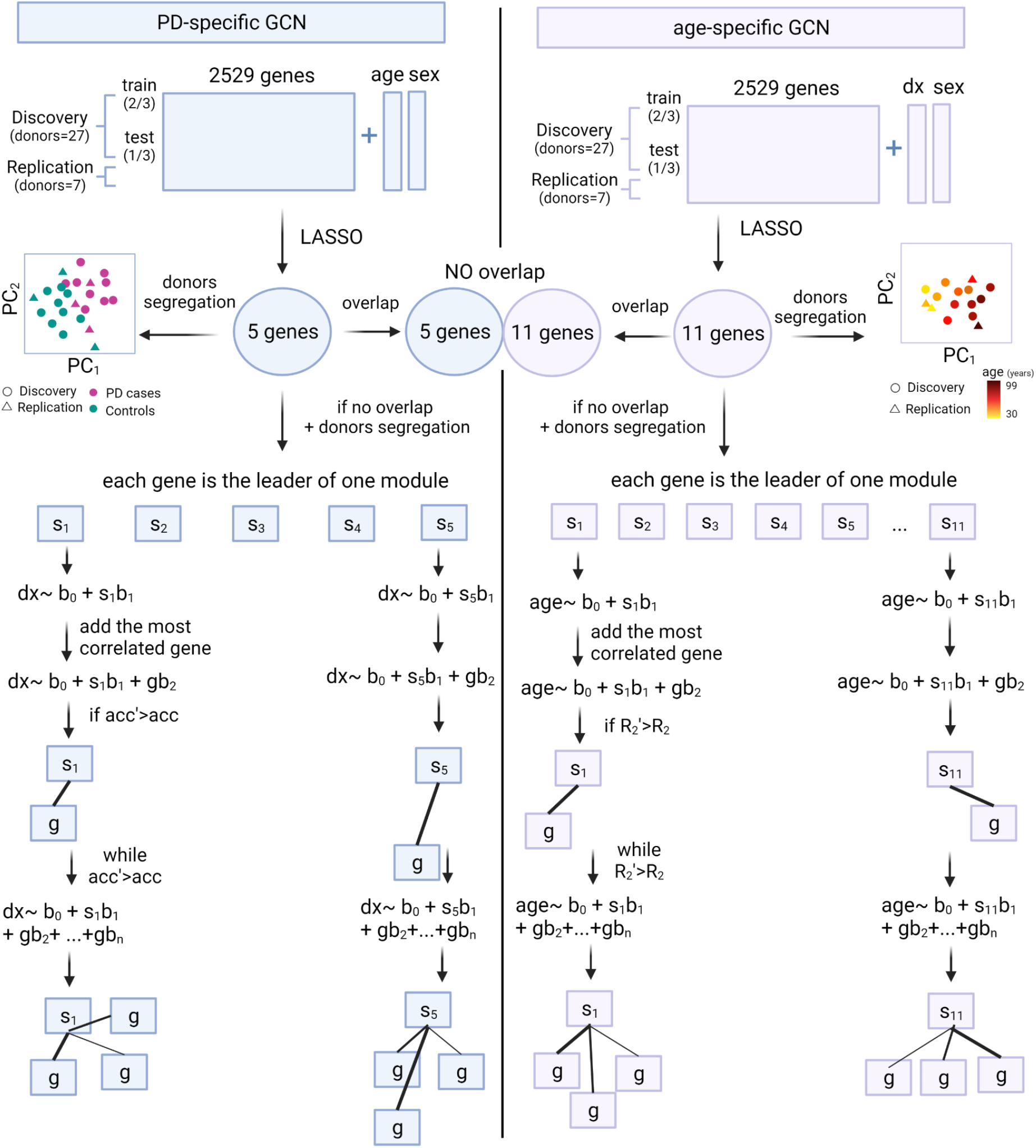
Pipeline for the creation of PD and age-specific GCNs from Oregon and Sepulveda datasets. We represent the creation process of each GCN across two columns (left for PD, right for age), starting from top to bottom. We started with the expression matrix of integrated single-cell samples from discovery and replication datasets, with additional columns for sex, age and diagnosis. We splitted the discovery samples into train (⅔) and test (⅓). The train samples were further splitted into 3 folds for cross-validation. We used the train set for selecting the most relevant genes for predicting age or PD separately using cv.glmnet with LASSO approach. We repeated the train and test split at least 10 times and the train split into three folds at least another 10 times. We selected the best model based on its performance on the test and replication sets. We then obtained the overlap between genes selected for predicting age and PD. As we found no overlap between the gene sets we then created the GCNs by using each selected gene as the leader of one module of the corresponding network. For each module, we kept adding the genes with highest correlation with the leader into the module, while the linear model R^2^ increases.

**Supplementary fig. 3.**
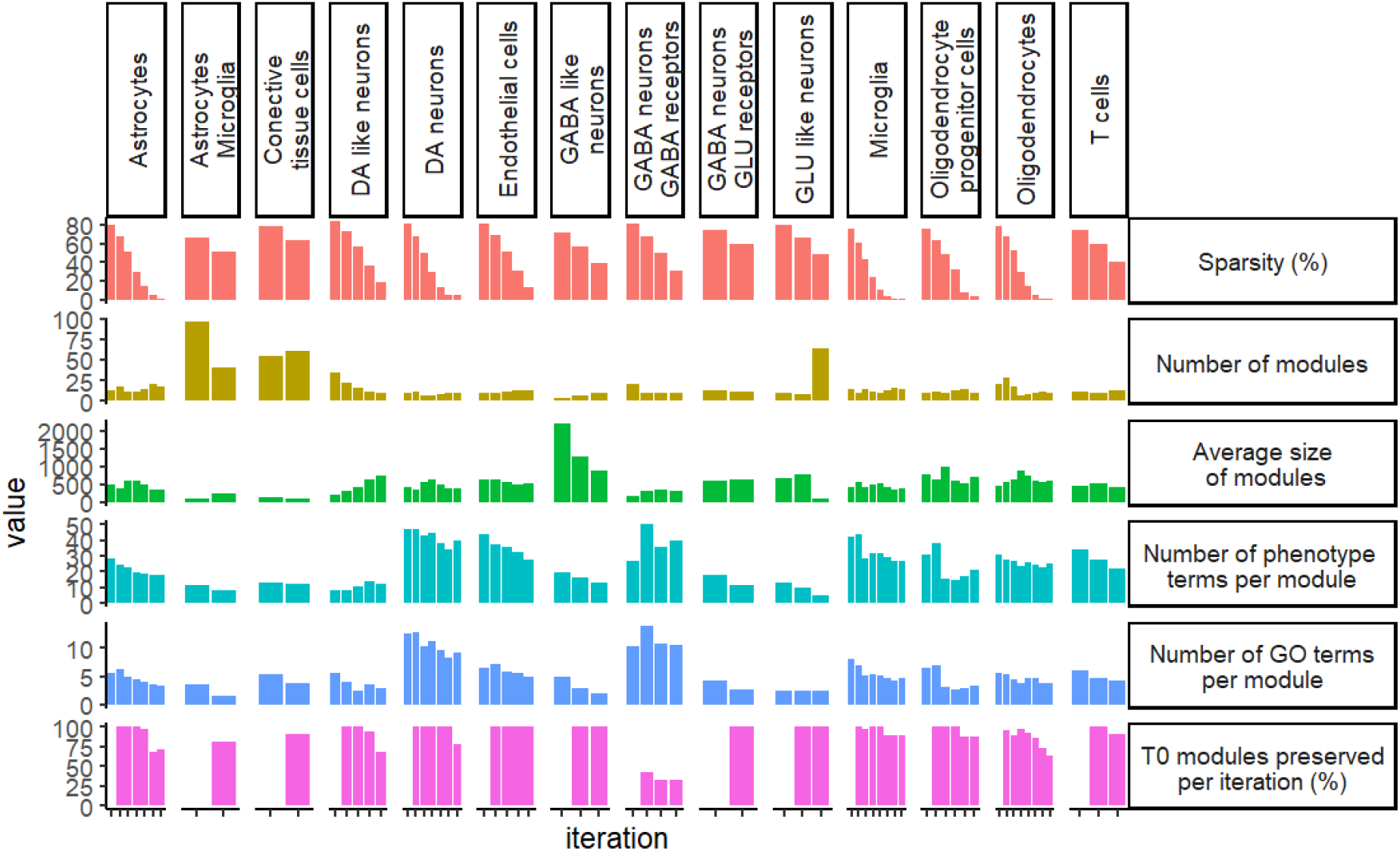
The creation of multiple scGCNs: reducing the sparsity while retaining the main features. Cell types within each plot column. X-axis of each plot refers to the iteration within which the scGCN was created from T_0_ to T_n_, where the maximum number of iterations is 6. Plots at rows correspond to the features observed from networks including sparsity (%), scGCNs main features and the % of T_0_ modules that are preserved in the rest of iterations based on Z summary press estimates.

**Supplementary fig. 4.**
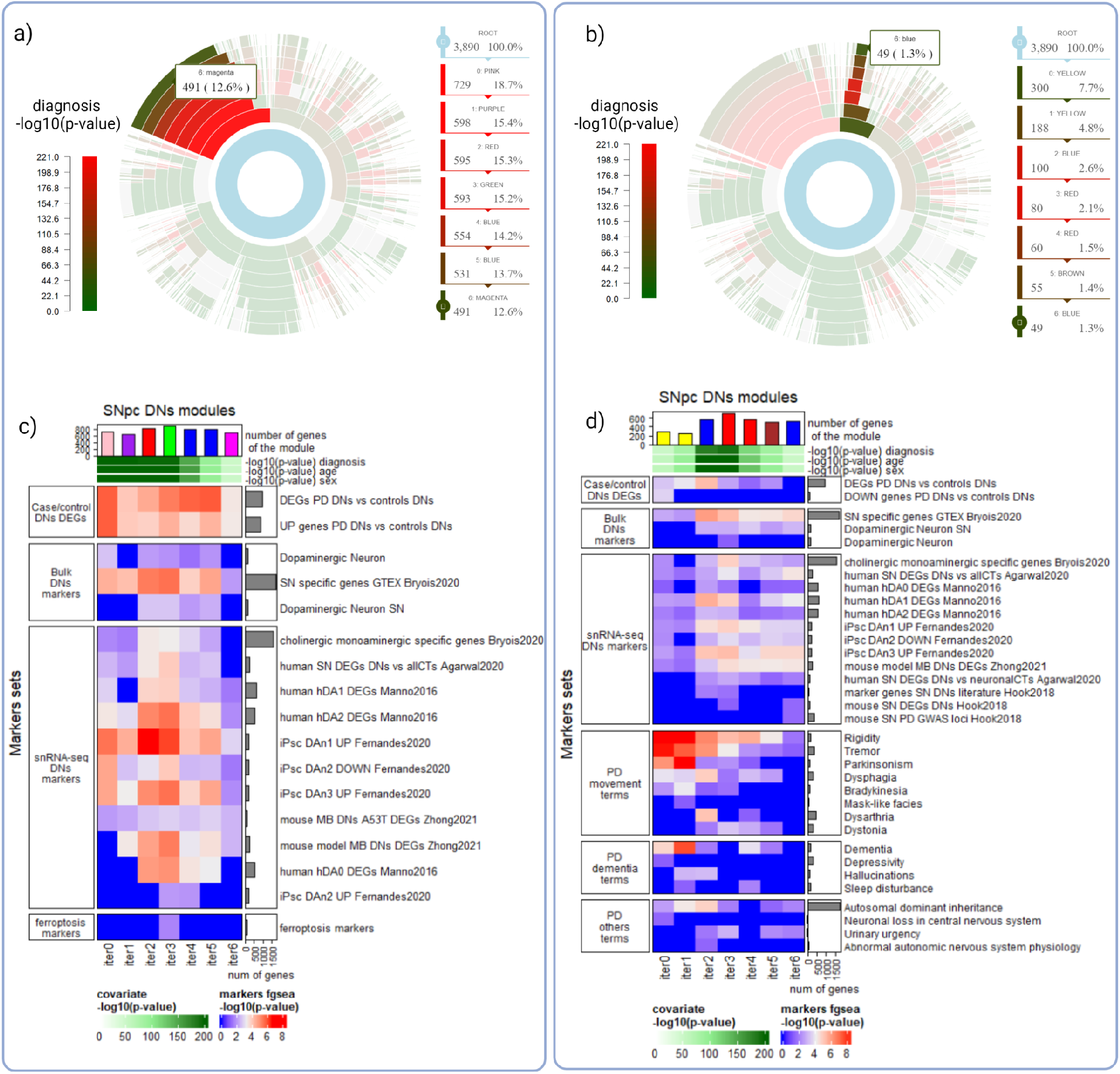
Study of scGCNs modules across iterations. Sunburst plots at **a)** and **b)** show as many rings as pseudo-cells iterations of DNs. The inner ring represents T_0_, the outer ring represents T_6_. All rings are divided into segments, one for each module of the corresponding scGCN. The size of the segment represents the number of genes within the module. M_1_ in T_0_ is highlighted at sunburst a). The majority of genes within M_1_ remain together across iterations. Colors at the segment show gene correlation with clinical diagnosis. At b) we see, highlighted, a particular gene module, M_2_, whose gene composition changes significantly across iterations just to get better association with clinical diagnosis at iterations 3 and 4. Heatmaps at **c)** and **d)** show, for M_1_ and M_2_, respectively, the different sets of annotations (at rows) as they evolve through iterations (at columns, from left to right). From top to bottom, we have characterized the DNs modules showing the number of genes that make up each of these modules (bars height across iterations) and their correlation with clinical diagnosis, age and sex covariates (in green). Then, we use four different groups of gene markers to further annotate the modules: (1) DNs DEGs between PD cases and controls, (2) cell type markers from bulk RNA-seq studies, (3) DNs markers obtained from sc/snRNA-seq studies and (4) Parkinson’s disease associated terms obtained from the Human Phenotype Ontology (see methods). Cells at the heatmap show results of the gene set enrichment analysis tests. Only marker lists with at least one significant test are shown (see methods for the complete list). These plots are generated by the scGCNs R package to help the analyst on studying all models.

